# Enzyme Activation of Triggerable Polymeric Lysosome Targeting Chimeras

**DOI:** 10.64898/2026.09.03.749226

**Authors:** Badal Singh, Priyanath Das, Aniket Majee, Adam Tajdi, Anirudh Deverajan, Allison Lomeli, Amelia Talluri, Ranit Dutta, S. Thayumanavan

## Abstract

Targeted protein degradation (TPD) has been highly effective for intracellular targets, but extending this approach to extracellular and membrane-bound proteins remains difficult because most extracellular TPD (eTPD) strategies depend on ligand-targeting receptors (LTRs) whose expression and recycling vary across tissues. Existing LTR-independent, multivalent platforms expand the scope of eTPDs by eliminating this dependence on LTRs and enabling cancer-selective designs. Layering in an additional degree of selectivity through endogenously activatable triggers, such as overexpressed enzymes, can enhance the tissue tropism of the LTR-independent platforms and widen therapeutic window. Here we report triggerable polymeric lysosome targeting chimeras (tPolyTACs), which combine antibody-defined targeting with locally triggered covalent capture. tPolyTACs conjugate monoclonal antibodies to phosphatase-cleavable substrates that, upon engagement of endogenous cell-surface phosphatases, unmask a reactive quinone methide electrophile, covalently trapping the target complex and driving its clathrin-mediated internalization and autolysosomal degradation. We show tPolyTAC-mediated degradation of the membrane proteins EGFR, PD-L1, and cMET, and demonstrate that para-substituted electrophiles outperform ortho-substituted analogues likely due to more favorable active-site positioning. These results establish tPolyTACs as a modular, covalent, enzyme-responsive platform that resolves the efficiency-selectivity trade-off limitations in extracellular degradation.

## 1. Introduction

The development of targeted protein degradation (TPD) technologies has revolutionized therapeutic strategies by enabling the elimination of previously “undruggable” proteins.^1,2,3^ While most TPD approaches have focused on intracellular targets, there is growing interest in degrading extracellular and membrane-bound proteins, which are often implicated in cancer, autoimmune, and inflammatory diseases.^4^ Current extracellular TPD (eTPD) strategies, such as LYTACs, KineTAC, and MoDE-As, rely heavily on ligand-targeting receptor (LTR)-mediated internalization. For example, LYTACs employ synthetic glycopeptide ligands that engage the cation-independent mannose-6-phosphate receptor to reroute cell-surface and secreted targets to the lysosome^5^, KineTACs repurpose the atypical, non-signaling chemokine receptor CXCR7 as an internalizing LTR via a cytokine-Fc fusion arm^6^, and MoDE-As use small-molecule ligands for the hepatocyte-restricted asialoglycoprotein receptor to achieve liver-selective degradation^7^.

To circumvent this dependence on internalizing receptors, we have previously developed two complementary LTR-independent strategies that show that simple multivalent engagement at the cell surface can drive lysosomal degradation without co-opting a dedicated lysosome-shuttling receptor. In one form of this polymeric lysosome targeting chimeras (PolyTACs) platform, a targeting antibody is paired with a ligand-decorated polymer that multivalently and non-covalently engages an ancillary membrane protein.^8^ The resultant antibody-defined specificity is high and the degradation efficiency is dependent on the avidity of the ligands against the ancillary protein target. In a second form of the PolyTACs platform, reactive electrophiles displayed on the polymer scaffold crosslink nearby cell-surface functionalities.^9^ This process has an inherently high apparent ‘avidity’, because of the covalent nature of the interaction. Notably, the non-covalent and covalent PolyTACs are always active, meaning their degrader function is solely dependent on expression of cell surface target protein and the ancillary proteins. Enhancing tissue tropism by introducing an additional layer of spatial control could help eliminate any potential trade-off between efficacy and selectivity. To this end, we propose an in-situ-activated covalent strategy that is switched on only upon processing of the polymeric functionalities by a cell-surface enzyme. Specifically, we designed triggerable PolyTACs (tPolyTACs), which leverage endogenous cell-surface phosphatases to unmask a reactive quinone methide intermediate (Scheme 1). This intermediate covalently modifies cell-surface proteins, driving autolysosomal trafficking of the proteins-of-interest (POIs) and their subsequent degradation.

**Scheme 1.**
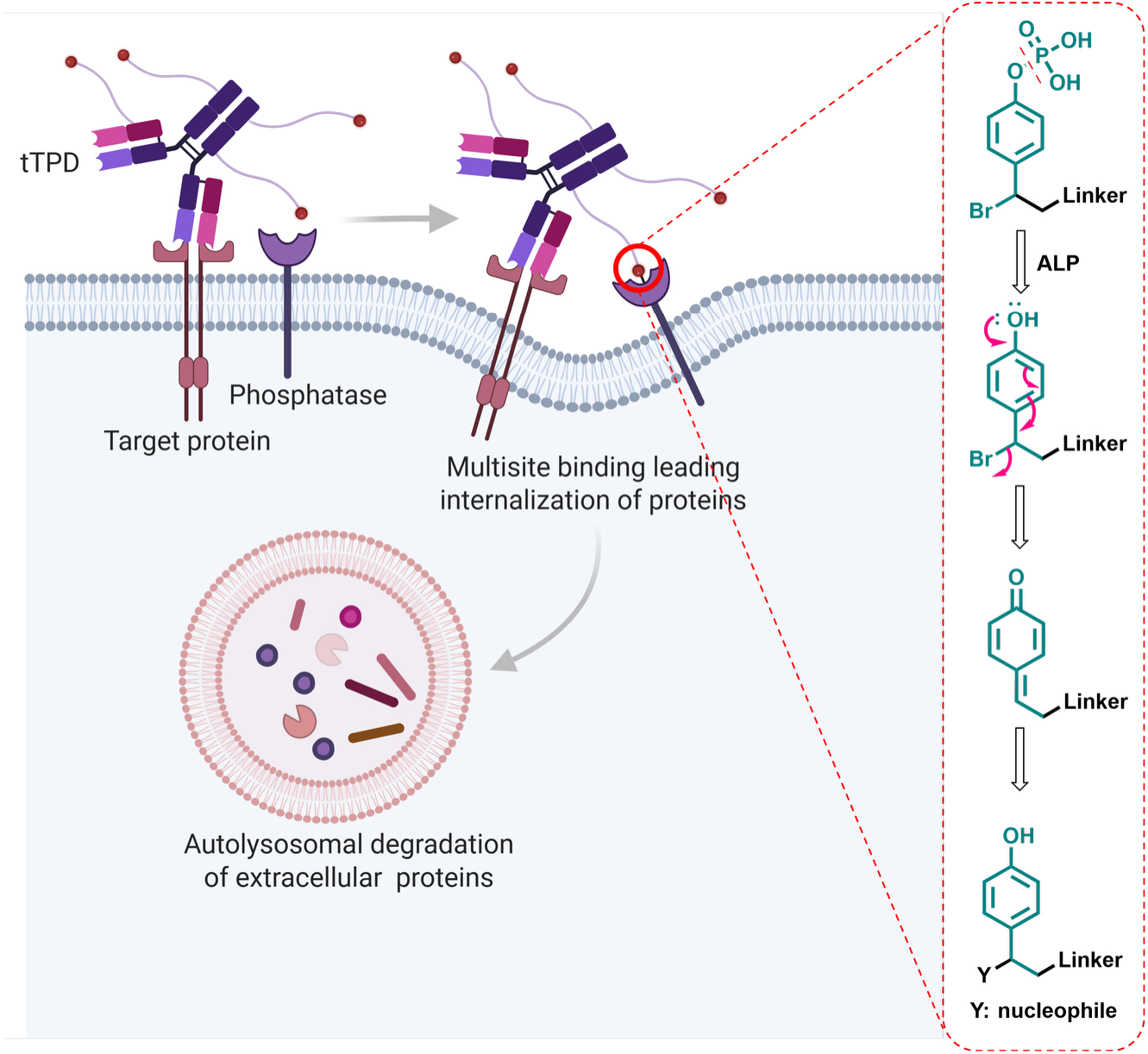
Multisite conjugation drives lysosomal degradation of a target protein. The tPolyTACs binds the target protein, and surrounding cell-surface phosphatases cleave its conjugated phosphate groups, unmasking a reactive electrophile that covalently engages the phosphatase active site (Y: active site nucleophilic amino acid). Multivalent covalent engagement traps the resulting complex at the cell surface, forcing its uptake by endocytosis; the endosome subsequently fuses with the lysosome, delivering the target protein for degradation.

The tPolyTACs are constructed via modular conjugation of monoclonal antibodies with polyethylene glycol (PEG) linkers bearing the enzyme-cleavable substrates. Upon enzymatic activation by cell surface phosphatases, the substrates generate quinone methide intermediates that covalently link nucleophilic residues on proteins. This allows durable surface tagging, plasma membrane invagination followed by cellular internalization (Scheme 1). We explore the structure-activity relationships of electrophile design, validate degradation in multiple disease-relevant targets, and extend the concept to soluble protein degradation. Overall, this strategy presents a flexible, covalent, and enzyme-guided framework for eTPD.

## 2. Results and Discussion

### 2.1. Design and synthesis of the tPolyTACs

The novelty and efficacy of this design rely on the dormancy of the highly reactive electrophile within a naturally stable aromatic functional group. We started by modifying a phosphatase-responsive phosphate-derived 2/4-hydroxybenzyl moiety that is substituted with a terminal alkyne for bio-orthogonal conjugation and a leaving group on the benzylic position. On the other hand, the antibody is modified with azido-PEG-NHS ester (MW 2000 Da) for Cu-assisted alkyne-azide click reaction with the phosphate derivative for tPolyTACs formation (Figure 1a).

**Figure 1.**
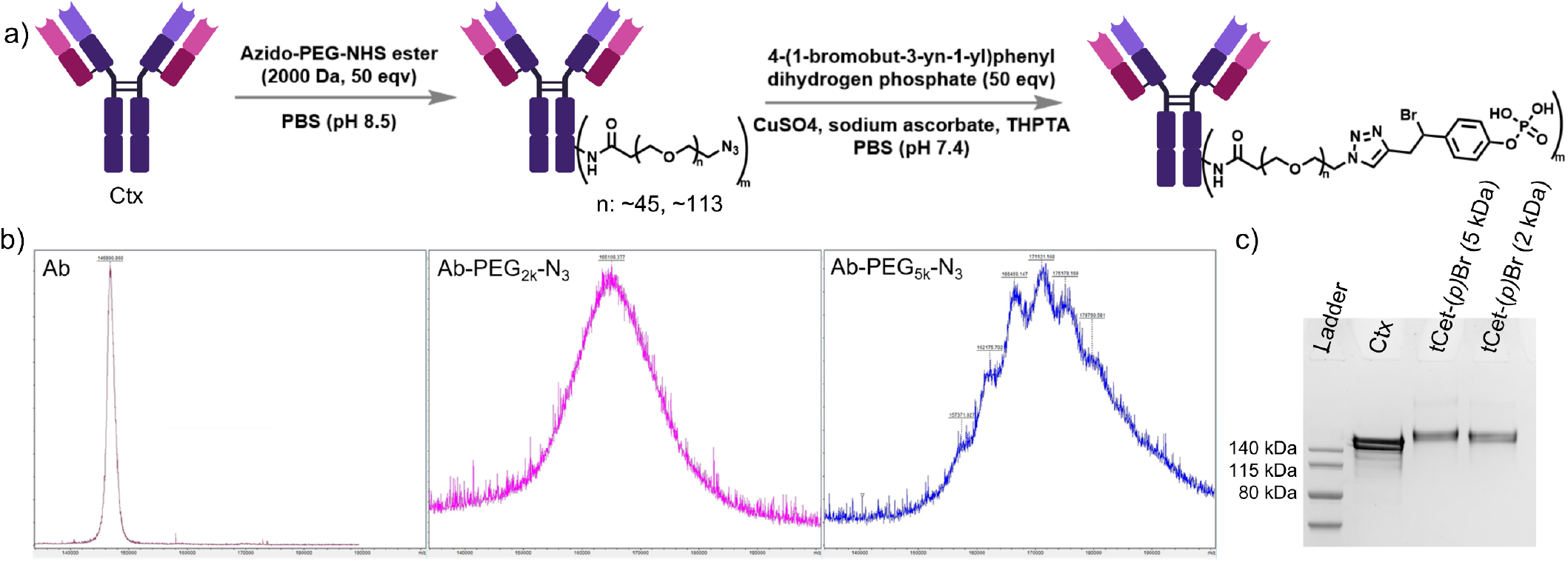
(a) Schematic illustration of preparation of tPolyTACs. (b) MALDI-MS analysis of long azido-PEG-NHS ester (MW 2 kDa and 5 kDa) conjugation to Ctx antibody (Ab). (c) SDS-PAGE gel showing final constructs, i.e., tCtx-(*p)*Br (2 and 5 kDa) indicating tPolyTACs with 2 and 5 kDa linkers respectively.

As the phosphatase-responsive substrate gets cleaved by transmembrane phosphatases and thereby transformed into a reactive Michael acceptor (a quinone methide derivative) in the phosphatase active site, it can react with a proximal nucleophilic functionality of an amino acid sidechain (*cis* modification) or react with amino acids from any other neighboring proteins (*trans* modification) or escape the active site and be hydrolyzed to the corresponding benzyl alcohol. The chances of the quinone methide to be hydrolyzed *vs cis* or *trans* covalent labeling would depend on the leaving group’s positioning and lability.^10^ For the optimized scenario, we expect covalent attachment of the phosphatase-responsive substate to cell surface proteins, especially phosphatases, to aid the degradation of membrane POI targeted by the antibody. Disappearance of the mAb band in SDS-PAGE is taken to indicate significant modification of the antibody. Number of azides per mAb was assessed using MALDI-MS, while the antibody consumption via Cu-assisted azide-alkyne cycloaddition to achieve the final tPolyTAC construct was characterized via SDS-PAGE (Figure 1c).

### 2.2. Determining the degradation efficacy using tPolyTACs and its structural optimization

We realized from previous reports that epidermal growth factor receptor (EGFR) proteins are surrounded by certain cell surface phosphatases.^11^ This convinced us to develop a phosphatase-assisted eTPD of EGFR. For EGFR targeting, we chose the therapeutic EGF-EGFR-binding antibody cetuximab (Ctx)^12^ and modified it with a phosphatase-substrate bearing a *para*-benzyl bromide functional group, yielding the degrader tCet-(*p*)Br. As we treated the EGFR-expressing MDA-MB-231 cell line with the tCet-(*p*)Br for 24 h, we observed efficient degradation of the POI, EGFR (Figure 2a,b). This supports our hypothesis of utilizing activatable covalent bond formation with non-LTR proteins like phosphatases to initiate lysosomal trapping of any extracellular POI for degradation. We also performed flow cytometry and confocal imaging using dye-labeled antibodies with complementary binding epitope to Ctx. Compared to the controls, we observed significant depletion in POI expressions after treating the tPolyTACs for 24 h in MDA-MB-231 cells.

**Figure 2.**
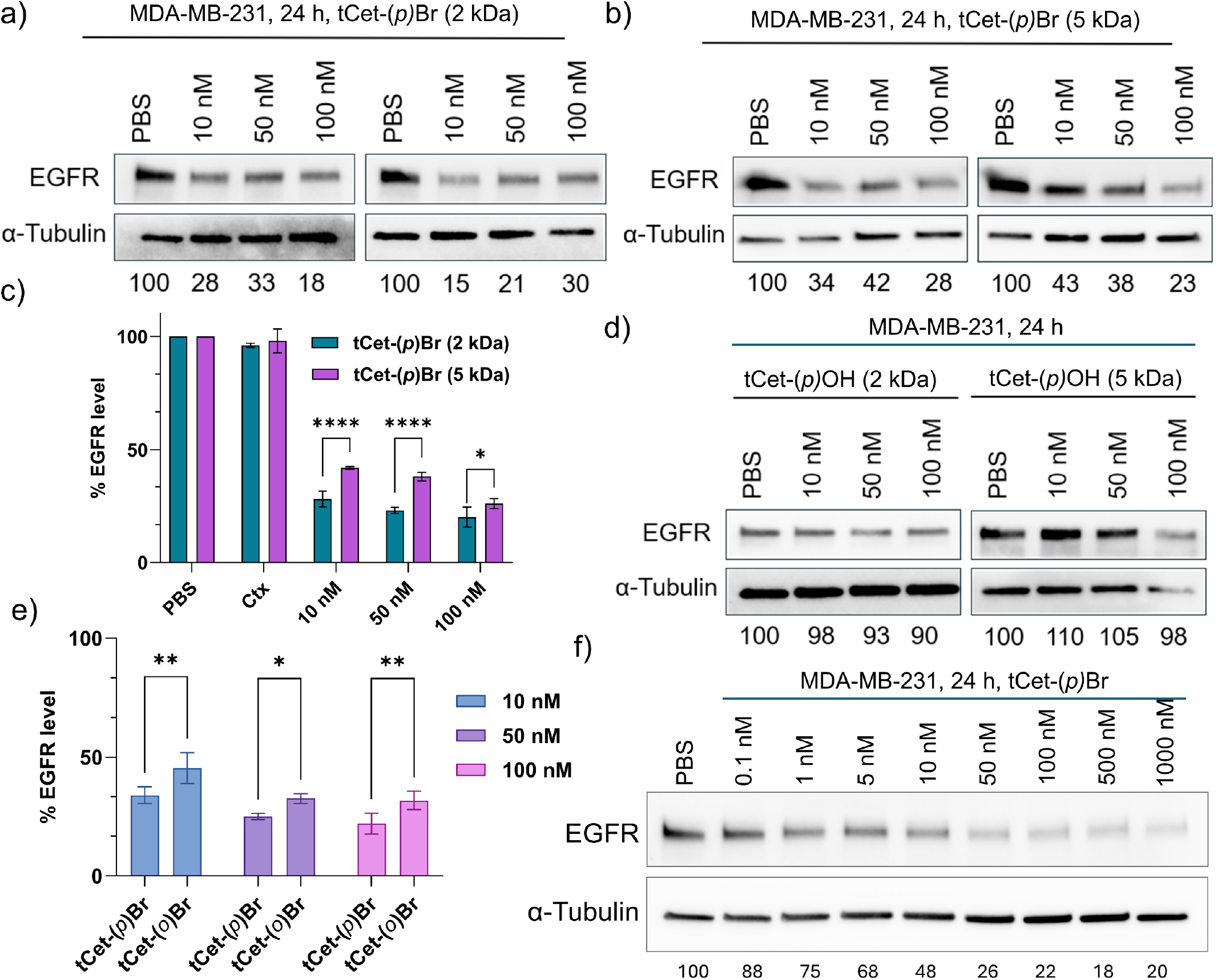
(a,b) Western blot results showing degradation of EGFR due to tCet-(*p)*Br (2 and 5 kDa respectively) treatment in MDA-MB-231 cells for 24 h. (c) Comparative flow cytometry results showing reduced cell surface expression of EGFR due to treatment of tCet-(*p)*Br (2 and 5 kDa). Results were analyzed by two-way ANOVA. (d) Western blot results showing degradation of EGFR due to tCet-(*p)*OH (2 and 5 kDa) treatment in MDA-MB-231 cells for 24 h. (e) Flow cytometry results showing degradation of EGFR due to tCet-(*o)*Br and tCet-(*p)*Br treatment in MDA-MB-231 cells for 24 h. Results were analyzed by two-way ANOVA. (f) Western blot results showing concentration-dependent EGFR degradation due to tCet-(*p)*Br treatment in MDA-MB-231 cells for 24 h.

We envision that, with proper optimization, the DC_50_ and D_max_ values can be improved in comparison to the current eTPD platforms. The first set of optimization is done by varying PEG linker length. We replaced the azido-PEG-NHS ester of 2000 Da molecular weight with another chain length of molecular weight 5000 Da. Since we noticed a small decrease in degradation efficacy, we opted for the shorter linker to avoid significantly affecting mAb binding affinity. Comparative flow cytometry confirmed that both the 2 kDa and 5 kDa linker constructs produced significant, dose-dependent reductions in cell-surface EGFR relative to PBS and Ctx controls, with the shorter linker giving modestly better degradation at each concentration tested (Figure 2c). The non-triggerable control PolyTAC tCet-(*p*)OH analogue, which lacks the electrophilic leaving group, showed little to no degradation of EGFR at either linker length, confirming that covalent bond formation, rather than mAb binding alone, drives internalization and degradation (Figure 2d). Also, the para-substituted tPolyTAC outperformed its ortho-substituted counterpart at every dose tested, and degradation remained concentration-dependent from 0.1 nM to 1 µM (Figure 2e,f). This difference likely reflects the relative positioning of the electrophilic center for nucleophilic attack: in the para-substituted isomer, the electrophile projects away from the linker and is well exposed to nucleophilic active-site residues, whereas in the ortho-substituted isomer the electrophile sits closer to the linker attachment point and may be sterically shielded by it, or held in a geometry that is poorly aligned for nucleophilic addition within the phosphatase active site^10^.

### 2.3. Checking modularity for different target proteins

We investigated the broad applicability of the system by targeting two other POIs, *viz*. PD-L1 and cMET. cMET dysregulation drives oncogenic signaling that fuels tumor cell proliferation, invasion, and resistance to EGFR-targeted therapies.^13^ PD-L1 overexpression enables tumors to suppress T-cell activity and evade immune surveillance, sustaining disease progression.^14^ PD-L1 and cMET proteins are significantly expressed in MDA-MB-231 cells (Figure 3b,d). To prepare the corresponding tPolyTACs, *viz*. tAtz-(*p*)Br and tAmz-(*p*)Br, we modified the respective mAbs, namely Atezolizumab (Atz) and Emibetuzumab (Emz). Both tPolyTACs, upon treatment in MDA-MB-231 cells, showed efficient degradation of PD-L1 and cMET respectively (Figure 3b,d).

**Figure 3.**
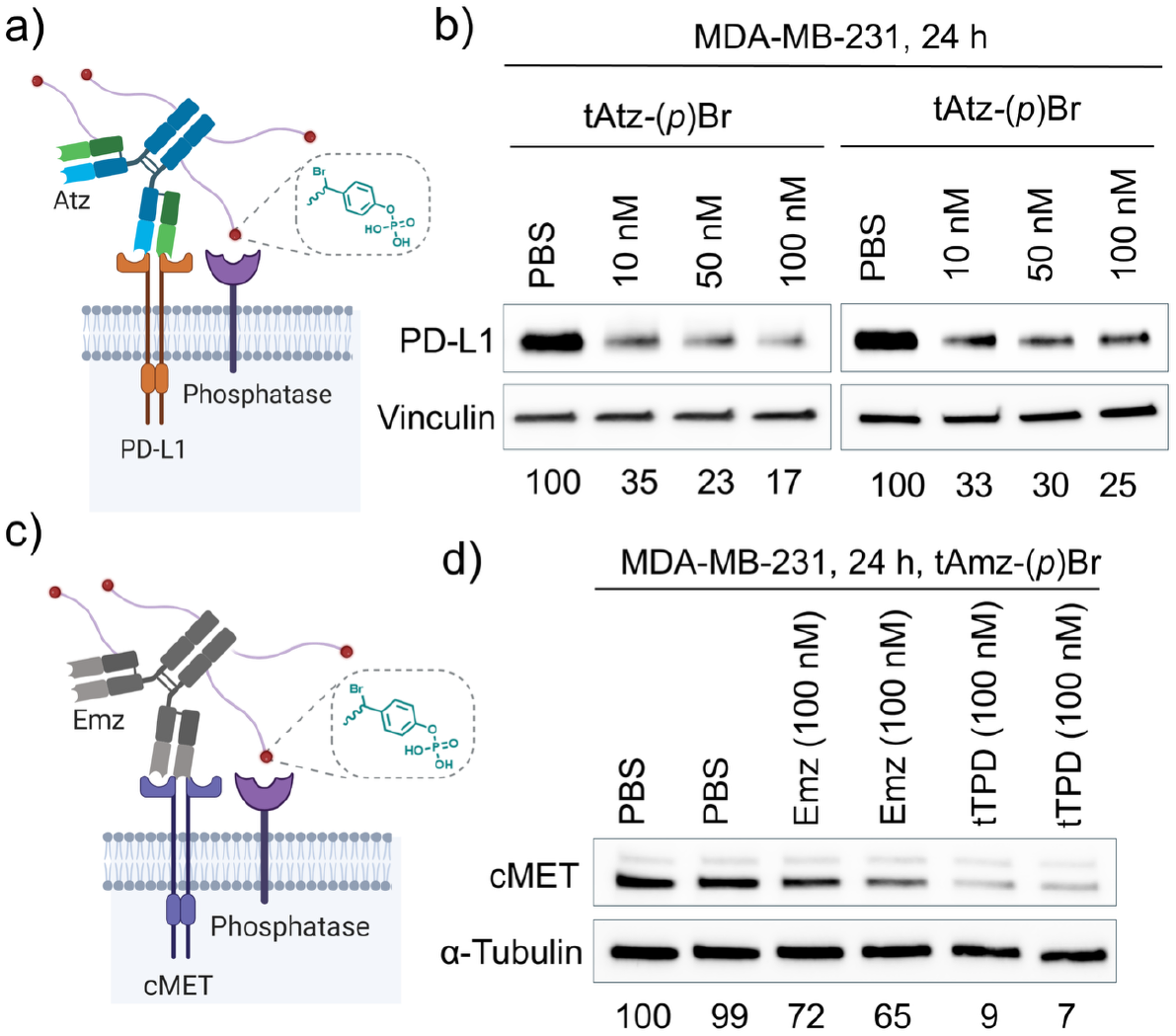
(a) Schematic illustration of PD-L1 targeting tAtz-(*p*)Br. (b) Western blot results showing degradation of PD-L1 due to tAtz-(*p*)Br treatment in MDA-MB-231 cells for 24 h. (c) Schematic illustration of cMET targeting tAmz-(*p*)Br. (d) Western blot results showing degradation of cMET by tAmz-(*p*)Br treatment in MDA-MB-231 cells for 24 h.

### 2.4. Mechanism of POI degradation

First, we investigated the importance of phosphatase active-site binding and responsiveness, the necessity of the leaving group, and the requirement for Ctx-EGFR binding. We used different formulations including: (i) tCet- (*p*)Br as positive control, (ii) tCet-(*p*)Br cotreated with Matuzumab that binds to a different epitope of EGFR, that is non-competitive to Ctx, (iii) tCet-(*p*)Br cotreated with Na_3_VO_4_ that competes with phosphate for the phosphatase active site, (iv) tCet-(*p*)Br cotreated with the activatable molecule itself, conjugated to azido-PEG-Ctx, namely (*p*)Br-PhP (i.e., *p*-bromomethyl)phenyl dihydrogen phosphate), and (v) tCet-(*p*)Br cotreated with the unreactive control tCet-(*p*)OH that lacks the leaving group. The inhibition of degradation in cases (*iii*) and (*iv*) shows the importance of phosphatase-activation of the Michael acceptor for degradation, and no inhibition of degradation in case (*v*) indicates the necessity of covalent blocking of the phosphatase active site (Figure 4a). To define the endocytic route responsible for internalization, cells were pretreated with chlorpromazine (CPZ, a clathrinmediated endocytosis inhibitor), methyl-β-cyclodextrin (MβCD, which disrupts cholesterol/caveolae-dependent uptake), EIPA (a macropinocytosis inhibitor), or cytochalasin D (CytoD, an actin polymerization inhibitor) prior to tCet-(*p*)Br treatment.^15^ CPZ nearly fully rescued EGFR levels, whereas MβCD, EIPA, and CytoD produced only partial rescue, implicating clathrin-mediated endocytosis as the predominant route of tPolyTAC-EGFR complex internalization (Figure 4b).

**Figure 4.**
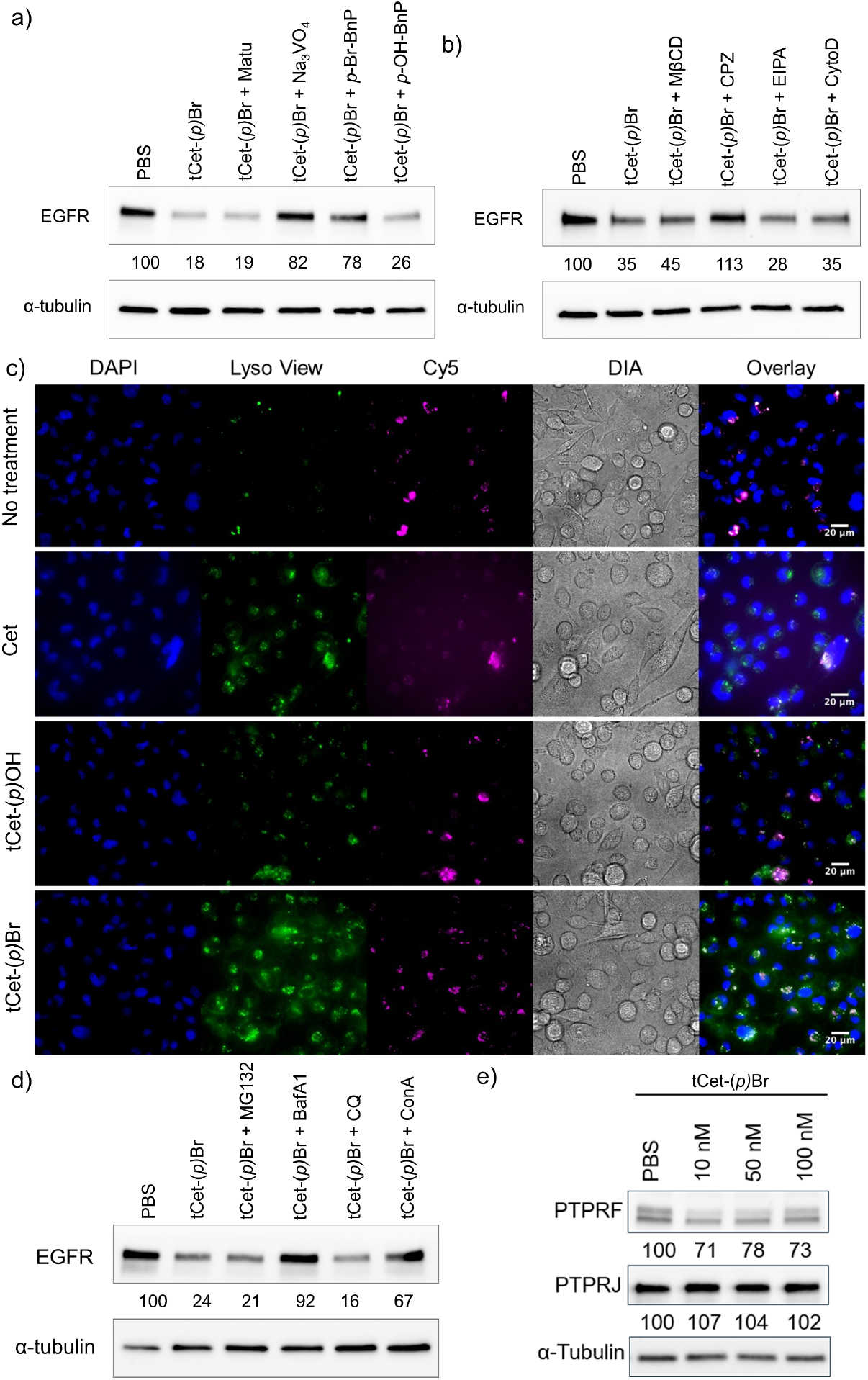
(a) Competitive experiments of tCet-(*p*)Br with phosphatase-active site blocking reagents showing the necessity of effective binding with Ctx and phosphatase-assisted electrophile generation in degrading Ctx. Inhibitors used include Matuzumab (Matu) that binds to EGFR and small molecules that can non-covalently or covalently target phosphatase active site. (b) Western blot analysis of EGFR degradation: MDA-MB-231 cells were pretreated with 10 µM chlorpromazine (CPZ), 0.5 mM methyl-β-cyclodextrin (MβCD), 10 µM 5-(N-ethyl-N-isopropyl)-amiloride (EIPA) and 10 µM cytochalasin D (CytoD) respectively for 15 min followed by washing and treatment of tPolyTAC for 24 h. (c) Confocal microscopy images of MDA-MB-231 cells after treating tCet-(*p*)Br (100 nM) for 24 h. Colocalization of LysoView and Cy5-labeled tCet-(*p*)Br indicates lysosomal trafficking of tPolyTAC and target proteins complex. (d) Western blot analysis of EGFR degradation: MDA-MB-231 cells were cotreated with proteasomal inhibitor (5 µM of MG-132) endolysosomal degradation inhibitors (100 nM BafA1 and 100 µM CQ) and autolysosomal degradation inhibitor (100 nM ConcA) and tCet-(*p*)Br for 24 h. (e) Western blot results showing PTPRF and PTPRJ degradation due to tCet-(*p*)Br treatment in MDA-MB-231 cells for 24 h.

Understanding the cellular fate of protein-tPolyTAC complexes is crucial to establish the mechanism of degradation and to further optimize the platform for therapeutic applications. Specifically, we sought to investigate whether the degradation of POIs by tPolyTAC is routed through the endolysosomal or autolysosomal pathway. First, we performed co-localization studies using confocal microscopy in MDA-MB-231 cells treated with fluorescently labeled tPolyTAC targeting EGFR, i.e., tCet-(*p*)Br and soluble IgG. Co-staining with organelle-specific markers revealed that the internalized protein-tPolyTAC complexes strongly colocalized with LysoView, indicating lysosomal trafficking of the POI (Figure 4c). Then we treated cells with bafilomycin A1, an inhibitor of lysosomal acidification, chloroquine (CQ, an endolysosomal acidification inhibitor), and ConA, an autolysosomal acidification inhibitor^16,17^, followed by tCet-(*p*)Br treatment. Bafilomycin A1 markedly inhibited POI degradation, while CQ had negligible effect on degradation efficiency, enforcing the conclusion that lysosomal processing is not dependent on endolysosomal trafficking (Figure 4d). However, because ConA treatment also inhibited degradation, POI turnover appears to proceed predominantly through the autolysosomal compartment. Co-treatment with the proteasomal inhibitor MG-132 similarly failed to rescue EGFR levels, arguing against a major contribution from proteasomal degradation (Figure 4d). Because phosphatase-responsive tPolyTACs act through neighboring cell-surface phosphatases, we also examined whether the receptor-type tyrosine phosphatases PTPRF and PTPRJ were themselves affected. tCet-(*p*)Br treatment modestly reduced PTPRF levels in a dose-dependent manner but left PTPRJ levels essentially unchanged, suggesting some selectivity among the phosphatase isoforms engaged by the electrophile (Figure 4e). This selective engagement is consistent with proximity-labeling proteomics studies that identify PTPRF, but not PTPRJ, as a component of the EGFR interactome^11,18^.

To conclude, these data indicate that tPolyTAC-mediated degradation occurs through the autolysosomal route, facilitated by covalent anchoring at the membrane. This mechanistic insight distinguishes tPolyTAC from ligandreceptor-based degraders and provides a rationale for further exploiting their unique trafficking behavior for targeted extracellular protein depletion.

### 2.5. Soluble extracellular protein degradation

While initial development of tPolyTACs focused on membrane-associated proteins, we next sought to explore their ability to degrade soluble extracellular proteins, which pose a unique challenge due to their diffusible nature and lack of cell anchoring. Soluble proteins such as vascular endothelial growth factor (VEGF), transforming growth factor beta (TGF-β), and immunoglobulin G (IgG) are key modulators of tumor progression, immune regulation, and inflammation.^19-21^ Strategies for their depletion could have broad therapeutic implications, particularly in cancer and autoimmune diseases. As an initial proof of concept for this class of diffusible target, we focused on IgG.

We therefore prepared tPolyTACs conjugated to monoclonal antibodies targeting IgG (anti-rabbit Fc), namely IgG-(*p*)Br. To visualize internalization and degradation, we performed confocal imaging using fluorescently labelled IgG and anti-IgG antibody decorated with the phosphatase-activatable groups. Post-treatment, we observed intracellular puncta colocalizing with LysoView, indicating successful uptake and trafficking to degradative compartments (Figure 5). These results demonstrate that tPolyTACs are effective degraders of not only membrane-associated proteins but also diffusible soluble extracellular proteins, provided the system is activated in proximity to cell-surface phosphatases. This expands the scope of tPolyTACs beyond surface-bound targets and opens avenues for neutralizing soluble pathological mediators in a targeted and controllable manner.

**Figure 5.**
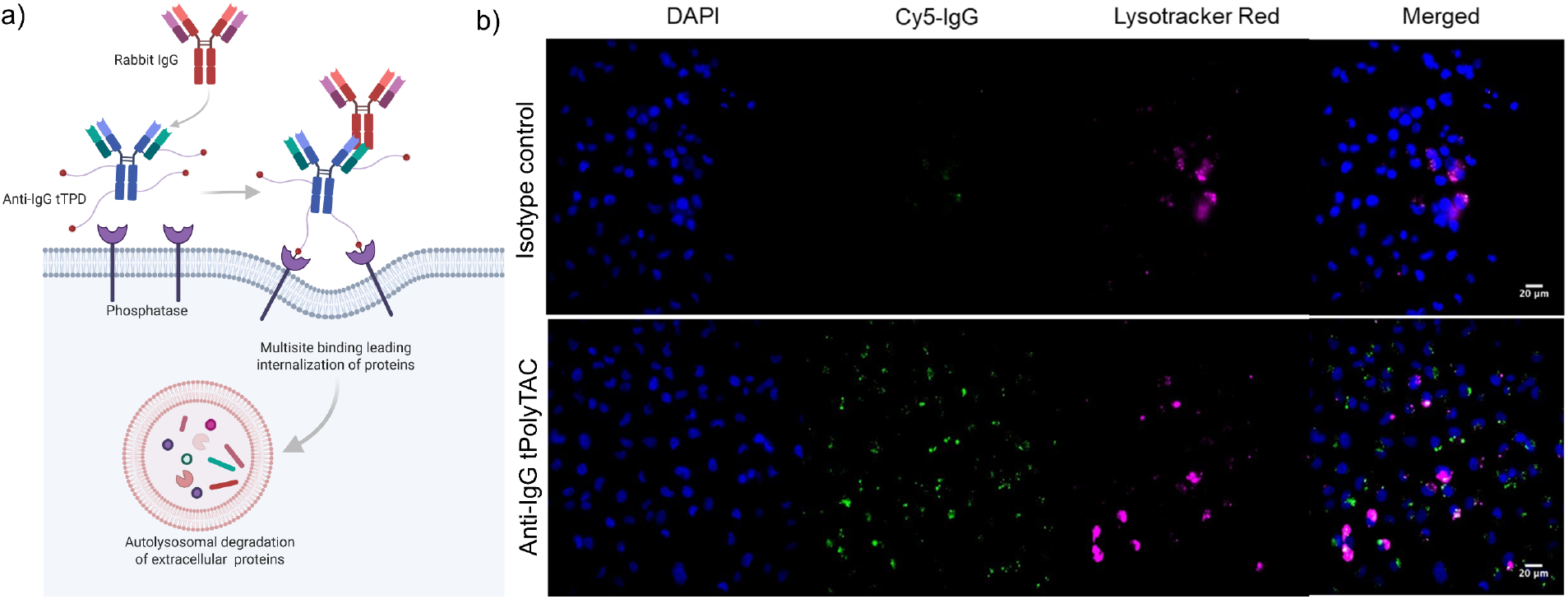
(a) Schematic illustration showing endolysosomal entrapment of extracellular soluble IgG due to the tPolyTAC IgG-(*p*)Br treatment. (b) Confocal microscopy images of MDA-MB-231 cells after treating Rb IgG and anti-Rb IgG-(*p*)Br (100 nM) for 24 h. Colocalization of LysoView and Cy5-labeled Rb IgG indicates lysosomal trafficking of IgG and the tPolyTAC complex.

## 3. Conclusion

In this study, we introduce tPolyTACs as a versatile and in situ activatable platform for targeted degradation of extracellular proteins, expanding the scope of current eTPD technologies. By leveraging cell-surface phosphatases as model activating enzymes to unmask reactive quinone methide intermediates, tPolyTACs achieve in situ activation and covalent tagging of target proteins, triggering their internalization and lysosomal degradation. Key findings include: (i) tPolyTACs are built by conjugating monoclonal antibodies to phosphatase-cleavable substrates. Local phosphatase activity unmasks a reactive quinone methide electrophile that covalently tags the target complex at the cell surface; (ii) The platform is modular across membrane-bound targets, achieving efficient degradation of EGFR, PD-L1, and cMET, with fine control over potency through optimization of linker chemistry and substitution position; (iii) Leaving-group placement tunes reactivity: para-substituted electrophiles consistently outperform ortho-substituted analogues, likely due to more favorable positioning for nucleophilic attack within the phosphatase active site; (iv) tPolyTAC-POI complexes are internalized predominantly via clathrin-mediated endocytosis and degraded through the autolysosomal, rather than the proteasomal or purely endolysosomal pathway; (v) The platform extends beyond membrane-bound proteins to diffusible soluble targets, as demonstrated for IgG.

This LTR-independent, activatable membrane protein engagement strategy circumvents the need for receptor recycling and expands the degradable target landscape to include proteins previously considered inaccessible. By uniting antibody-defined selectivity with covalent, enzyme-triggered efficiency in a single modular construct, tPolyTACs directly resolve the potential efficiency-selectivity trade-off. In this work, phosphatase served as a model cell-surface enzyme to establish and validate the tPolyTAC platform and future studies will extend this enzyme-responsive strategy to other disease-relevant cell-surface enzymes to broaden its therapeutic scope. Together, these results position tPolyTACs as a promising and broadly applicable technology for therapeutic protein degradation at the cell surface and in the extracellular milieu.

## Acknowledgment

We thank NIBIB of the NIH for support (EB037006).

